# A conditional histone H3.3 mutation in mice orthologous to a driver of paediatric diffuse instrinsic pontine glioma

**DOI:** 10.1101/2022.11.25.518018

**Authors:** Jeffrey R. Mann, Michelle C. W. Tang, Lee H. Wong

## Abstract

Most paediatric diffuse intrinsic pontine gliomas carry a lysine- to methionine-27 (K27M) mutation in the histone variant H3.3 encoded by the ‘H3.3 histone A’ (*H3F3A*) gene. To establish a pre-clinical model of the disease we made a Cre/*loxP* conditional H3.3K27M mutation at the orthologous *H3f3a* locus in mice. Importantly, expression of the mutant transcript is under endogenous *H3f3a* regulatory control. This system is distinct from others in which H3.3M27 is ectopically expressed, thereby providing a resource for the development of pDIPG models with orthologous regulation of mutant H3.3. Mice in which expression of the mutant transcript was induced with a nestin-*cre* transgene developed as dwarfs in the presence of intact growth hormone signaling.

## INTRODUCTION

A high proportion of paediatric diffuse intrinsic pontine gliomas (pDIPGs) or paediatric high-grade gliomas of the brainstem are associated with a somatic point mutation causing a lysine-to methionine-27 (K27M) amino-acid substitution in the histone variant H3.3 encoded by *H3F3A*. The H3.3K27M mutation is almost exclusive to pDIPG and is usually accompanied by mutations in ‘transformation related protein 53’ (*TRP53*) (Sturm et al., 2012; Wu et al., 2012). The H3K27 residue is integral to the function of polycomb repressor complex 2 (PRC2), a genome-wide transcriptional repressor of lineage-specific genes. The presence of H3.3M27 is thought to disrupt H3K27 methylation which in turn disrupts PRC2 chromatin localization, leading to widespread transcriptional deregulation (Chan et al., 2013; Lewis et al., 2013).

Preclinical models of the disease would greatly benefit the understanding of disease aetiology and the development of therapeutic regimes, and preclinical models in mice have been described (Cordero et al., 2017; Hennika et al., 2017; Pathania et al., 2017). However, in these cases mutant H3.3M27 protein was introduced ectopically, raising the question of whether the gliomas produced are the best representations of the human disease. It is relevant that K27M mutations at the second gene encoding H3.3, ‘H3.3 histone B’ (*H3F3B*), have not been described in pDIPG. This gene, at least in mice, is differentially expressed relative to *H3F3A* (Bramlage et al., 1997; Tang et al., 2015), suggesting that the spatial, temporal, or level of expression of *H3F3A* is an important disease determinant. A second disadvantage of preclinical models relying on ectopic expression strategies is that vector delivery methods are labour intensive, requiring substantial manipulation of animals. To circumvent these issues we made a Cre/*loxP* H3.3K27M conditional mutation at the orthologous *H3f3a* locus in mice in which expression of the mutant transcript is under endogenous *H3f3a* regulation. This mouse line provides an important resource for the development of preclinical pDIPG models which have an aetiological profile matched as closely as possible with the human disease.

## RESULTS & DISCUSSION

The gene targeting strategy in mouse embryonic stem (ES) cells is described (Materials & Methods section, Figure 1). Nineteen percent of clones picked and expanded were correctly targeted as assessed in Southern blots. In targeted cells, only the wild-type H3.3K27 coding sequence (cds) was expressed, driven by the endogenous *H3f3a* promoter (conditional allele, designated CK27M). On exposure to Cre recombinase, the H3.3K27 cds and *neo* expression cassette were excised, bringing the H3.3M27 cds under *H3f3a* regulatory control (mutant allele, designated M27) (Figure 1A). The predicted shifts in band sizes with and without Cre-mediated recombination were seen in Southern blots (Figure 1B). Also, a switch in allelic expression from H3.3K27-to H3.3M27-encoding transcript was seen in G418-resistant versus -sensitive clones, respectively (Figure 2A, B). The faint band corresponding to mutant product in lane 2 for RTPCR-2 (Figure 2B) could be explained by low level transcriptional read-through from the *neo* cassette driven by the *Pgk1* promoter. Nevertheless, under these circumstances no translation of H3.3M27 protein would be expected. These assays showed the genetic modification functioned as designed. Following germ line transmission, a CK27M mouse line derived from clone no. 8 was used for all studies reported here.

**FIGURE 1.**
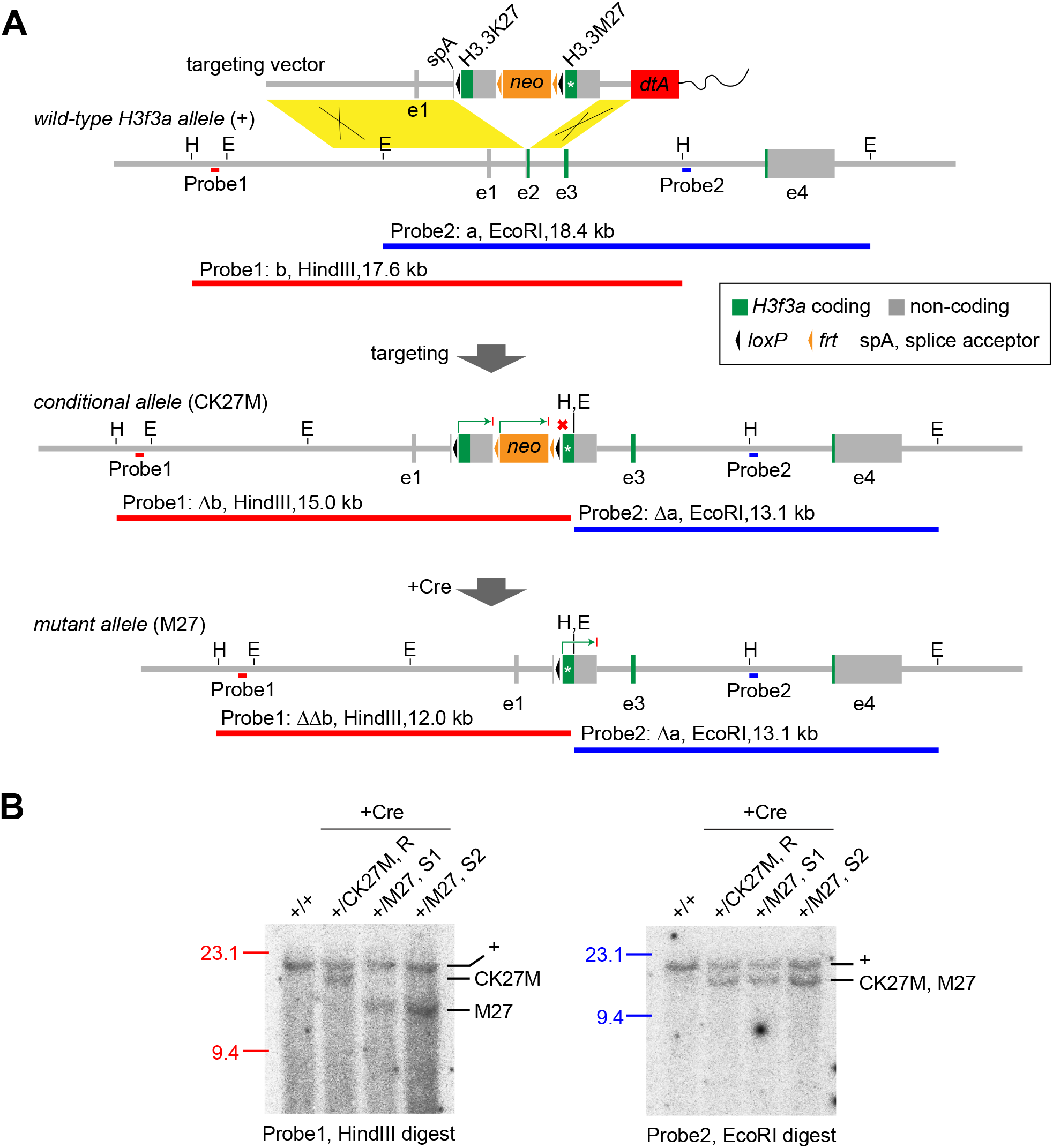
Gene targeting at *H3f3a*. (**A**) Design strategy and expected band sizes in Southern blots. E, EcoRI; H, HindIII. The promoter-less H3.3K27 and H3.3M27 minigenes/cassettes comprised the cDNA, the *H3f3a* UTR and the late-SV40.poly(A) signal sequence. The *neo* and *dtA* cds for positive and negative selection respectively were driven by the murine ‘phosphoglycerate kinase 1’ (*Pgk1*) promoter and terminated with the bovine GH poly(A). The targeting vector was linearized with XhoI. On targeting, the three internal cassettes replaced most of *H3f3a* exon 2 (e2) to bring the H3.3K27 cassette under endogenous *H3f3a* regulatory control (CK27M allele)—here the *neo* cassette can be removed with Flp recombinase if desired. On exposure to Cre (+Cre) the H3.3K27 and *neo* cassettes were excised, bringing the H3.3M27 cassette under regulatory control (M27 allele). Transcription (arrow), no transcription (cross). In the targeting vector, the H3.3M27 cds can be replaced with any other cds between unique BamHI and HindIII sites. (**B**) Southern blots of subclones resistant (R) and sensitive (S) to G418. Predicted band sizes were seen, demonstrating conservative recombination (clone #8). Primers used to make probes by PCR, 5′-3′: Probe 1, #1665, ACTA GAAA AACC GGCC AA, #1666, GCAT ATAC GGAG TCAG GGA, amplicon 306 bp. Probe 2, #894, ACCA TGCT GGGC TCTT TAC, #895, AACT GTGC TAGG CACA GC, amplicon 295 bp.

**FIGURE 2.**
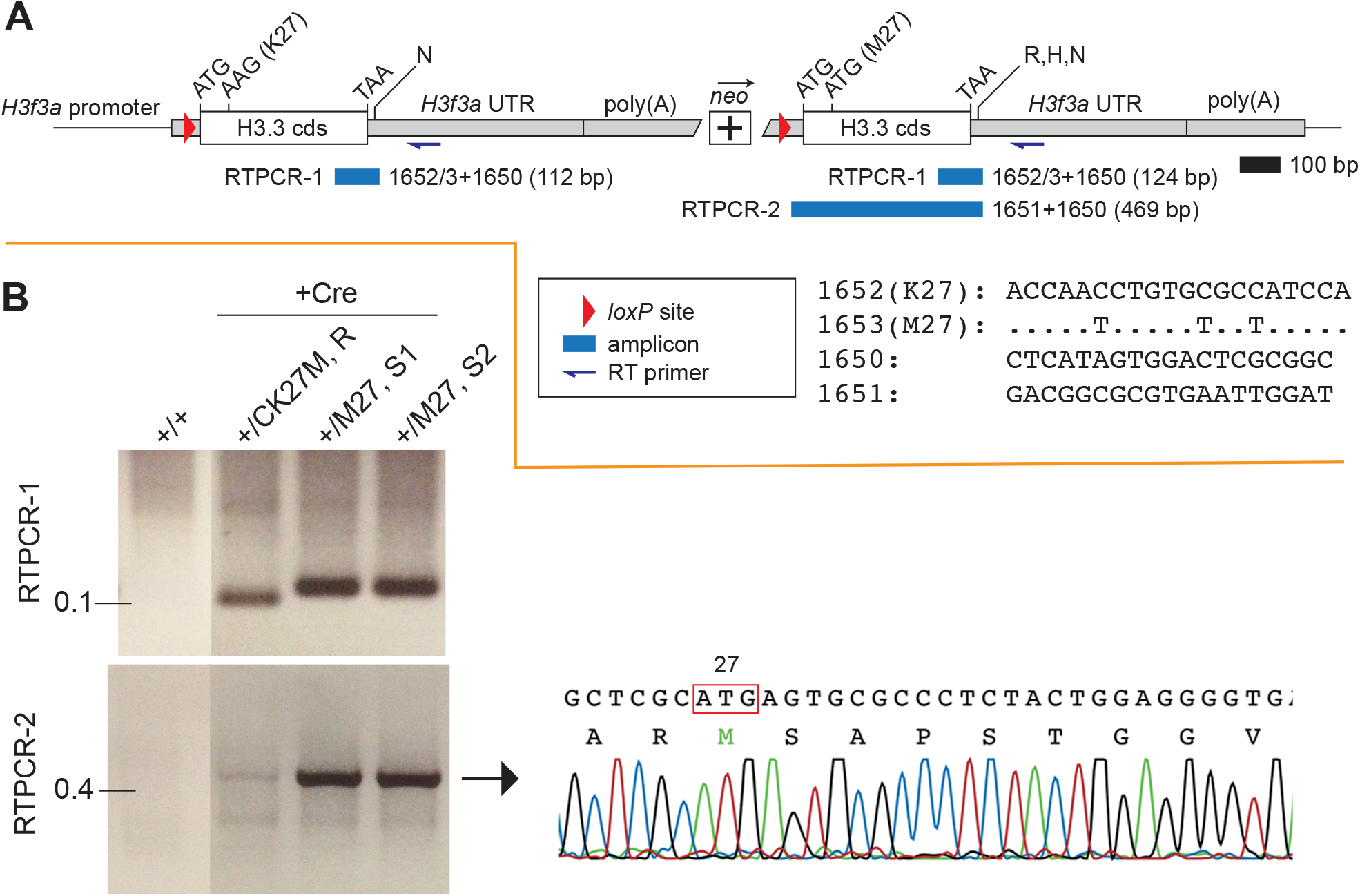
Expression of the CK27M allele in targeted ES cells before and after exposure to Cre recombinase. (**A**) Primers and location of expected RTPCR amplicons (blue bars) derived from the three internal expression cassettes—wild-type H3.3K27, *neo*, and mutant H3.3M27— targeted to exon 2 of the *H3f3a* locus. These primers are specific to the genetically modified allele and do not recognize endogenous sequences. In RTPCR-1, the three primers can amplify the H3.3K27-and H3.3M27-encoding transcripts in the same assay, with the former transcript being slightly smaller. In RTPCR-2, the two primers can amplify only the H3.3M27-encoding transcript. R, EcoRI; H, HindIII; N, NotI, restriction sites. (**B**) RTPCR assays showing the switch in expression in resistant (*neo* intact), to sensitive (*neo* excised), ES cell clones. RTPCR-1: switch from H3.3K27 transcript to H3.3M27 transcript. RTPCR-2: from faintly detectable expression to expression of the H3.3M27 transcript. +/+, wild-type ES cells; +/CK27M, G418-resistant (R) subclone; +/M27, G418-sensitive (S) subclone. Sequencing of the product in RTPCR-2 confirmed the presence of the M27 codon.

Homozygous CK27M mice were overtly normal and fertile. To see if inducing the mutation with Cre was capable of eliciting a biological effect in mice, +/+, 0/Tg-*nes*.*cre* females (mouse strain congenic B6) were mated to CK27M/CK27M males (mouse strain isogenic B6) to produce +/ CK27M, Tg-*nes*.*cre*/0 experimental and +/CK27M, 0/0 control pups in an expected 1:1 ratio. In experimental animals, extensive conversion of the CK27M to M27 allele was expected to occur in the developing central and peripheral nervous systems because of Cre expression in these tissues (Tronche et al., 1999). The presence of the mutant protein could not be confirmed because of the absence of a specific antibody. We avoided the use of epitope-tagged H3.3M27 in our system because of the high potential for artifacts: H3.3 is one of the most conserved proteins in the Metazoa—only histone H4 is more conserved. While tagged H3.3 incorporates into chromatin, normal functionality is questionable given possible adverse effects on nucleosome structure and tail post-translational modifications. For example, using gene targeting at *H3f3a* we constitutively tagged H3.3 with the MYC epitope. While ES cells were overtly normal, we were unable to obtain chimaeras after injecting several independent clones into blastocysts. This was a very unusual result obtained with the cell line used. Typically, our targeted clones give extensive chimaerism (J.R. Mann, unpublished data).

Experimental pups were of normal size at birth, but were less than 70% of normal weight by 8-10 days post-partum despite having fed normally—indicated by normal quantities of milk present in the stomach (Figure 3A). They remained small to early adulthood (Figure 3A) and throughout life. Both sexes were unable to produce young when mated to wild-type animals over a six month period. Over this time they remained in good health, then were culled. The growth phenotype was similar to proportional dwarfism seen for disruptions in the ‘growth hormone’ (GH) signaling pathway or somatotropic axis. However the urine of male adults contained normal concentrations of major urinary protein (MUP) (Figure 3B), a marker of GH signaling. GH activates liver ‘signal transducer and activator of transcription 5’ (STAT5), which then activates MUP (Norstedt and Palmiter, 1984; Waxman and Celenza, 2003).

**FIGURE 3.**
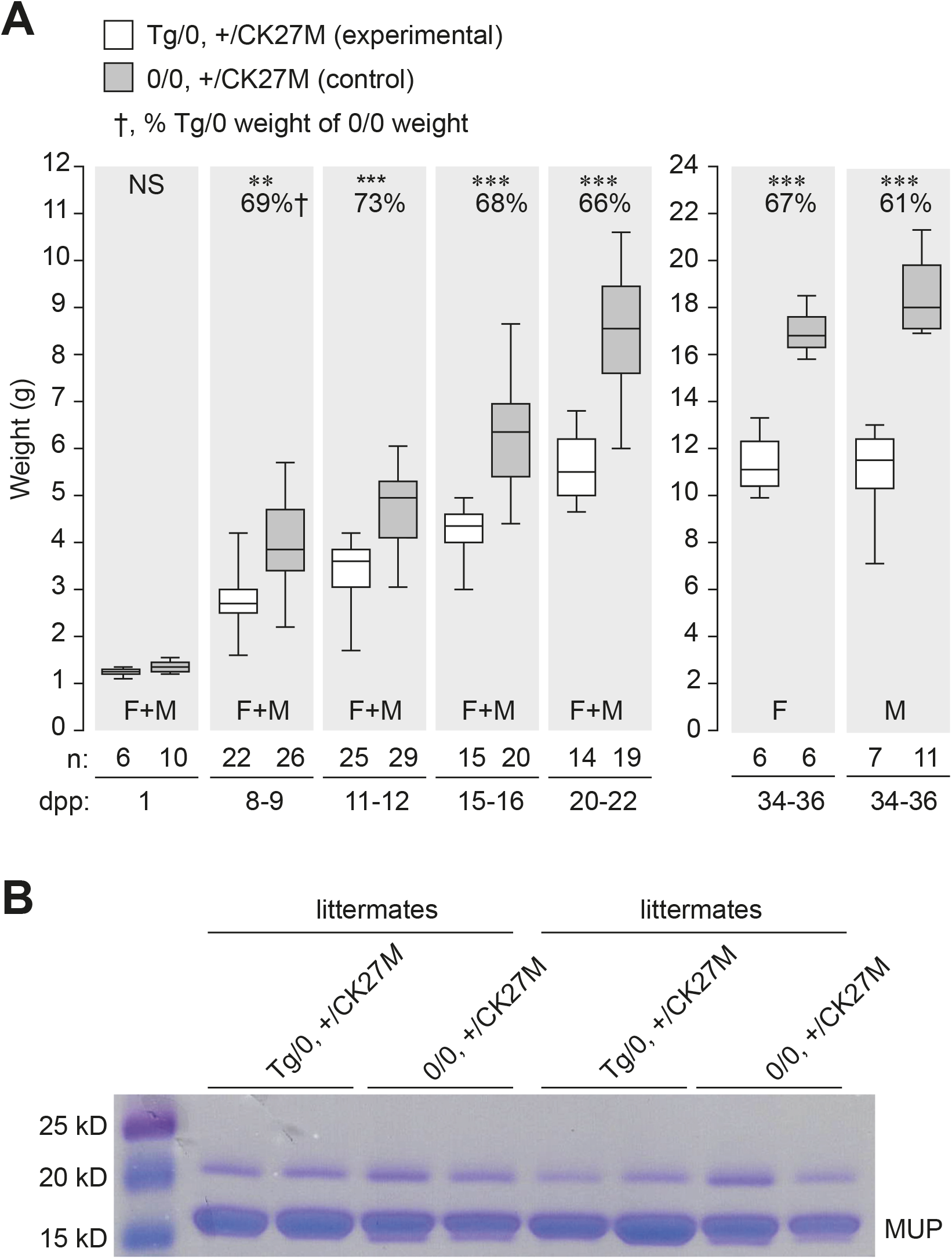
Assays of experimental and control animals born from +/+, 0/Tg-*nes*.*cre* ♀ *×* CK27M/CK27M, 0/0 ♂ matings. (**A**) Weight gain, displayed as box and whisker plots. Paired *t*-test, where one pair is one experimental and one control pup from the same litter: not significantly different (NS); significantly different at 0.05 (**) and 0.01 (***). F+M, weights in relation to sex were not distinguished in the data presented (1-22 days post partum or dpp). Female (F) and male (M) weights are plotted separately at 34-36 dpp. (**B**) Polyacrylamide gel electrophoresis of urine obtained from two sets of two experimental and two control adult male littermates at 12 and 18 weeks of age. Gels were stained with Coomassie Blue.

We have described a conditional Cre/*loxP* point mutation at the *H3f3a* locus in mice that encodes putative mutant H3.3M27 protein, a driver of pDIPG. Importantly, expression of the mutant transcript is under endogenous *H3f3a* regulatory control, and is therefore may be expressed like the orthologous human mutant transcript. This genetically modified mouse line provides a valuable resource for the development of preclinical models of pDIPG: first, endogenous *H3F3A*-directed expression of non-tagged H3.3M27 may be critical for developing a preclinical model which best represents the human disease, and second, the Cre/*loxP* system is relatively convenient for inducing the mutant protein at any stage of development. Also, the gene modification system described is easily adapted to produce other conditional point mutations in *H3F3A* which could serve as preclinical models of disease, such as H3.3G34R/V in non-brainstem gliomas and H3.3G34W/L in chondrosarcoma (Sturm et al., 2012; Behjati et al., 2013).

The survival of mice in which the mutant transcript would be induced in the central and peripheral nervous system of the early fetus was surprising given the widespread disruption of gene expression attributed to the mutant H3.3M27 protein (Chan et al., 2013; Lewis et al., 2013). The dominant-negative effect of dwarfism in mice with tissue-specific heterozygosity for the induced mutation cannot be accounted for by deficiency of wild-type H3.3 protein because even homozygous *H3f3a* mutants are close to normal size (Jang et al., 2015; Tang et al., 2015). These phenotypic effects of putative H3.3M27 protein require further investigation.

## MATERIALS & METHODS

The *H3f3a* gene targeting vector was constructed according to the Cre/*loxP* conditional allelic replacement system as described (Tang et al., 2013; Tang et al., 2015) and the full sequence deposited (Genbank accession: MH396437). The ES cell line Bruce7 of strain B6.PL-*Thy1*^a^/CyJ (B6) (Jackson Lab. stock. no. 000406) (Abbondanzo et al., 1993) was used for gene targeting. Recombinant ES cells were injected into strain BALB/c blastocysts for chimaera production. Male chimaeras were mated to strain B6 females for germ line transmission. PCR primers used to make probes for Southern blots as described in Figure 1: 5′-3′: Probe 1; #1665, ACTA GAAA AACC GGCC AA; #1666, GCAT ATAC GGAG TCAG GGA; amplicon, 306 bp. Probe 2; #894, ACCA TGCT GGGC TCTT TAC; #895, AACT GTGC TAGG CACA GC; amplicon, 295 bp.

ES cells heterozygous for the conditional allele were electroporated with the pCAGGS-*cre* plasmid (Araki et al., 1997) (10^7^ cells, 20 μg of circular vector) and plated (1000 cells/10 cm plate). Single colonies were picked and replica-plated into medium plus and minus G418 to obtain resistant and sensitive subclones. The former did not take up the plasmid or failed to undergo recombination, while the latter were putative Cre-mediated recombinants. These clones were used to assay expression using RTPCR and recombination using Southern blots. Total RNA was extracted from ES cells using the High Pure RNA Isolation Kit (Roche Diagnostics) and RTPCR performed using the High Capacity cDNA Reverse Transcription Kit (Applied Biosystems). Southerns blots were performed as described (Sambrook et al., 1989).

The CK27M allele (*neo* cassette present) was bred to homozygosity to produce a homozygous line. Subsequent to the studies reported here, this line has been bred to be also homozygous for a conditional null allele of *Trp53* (Jackson Lab. stock no. 008642) (Marino et al. 2000) and a conditional β-galactosidase reporter (Jackson Lab. stock no. 019101) (Soriano, 1999). The B6.Cg-Tg(Nes-cre)1Kln/J transgene (Jackson Lab. stock no. 003771) (Tronche et al., 1999) was used for expressing Cre in mice. The nestin sequences drive *cre* in the neuroepithelium through an enhancer in intron-2 (Zimmerman et al., 1994). This transgene is abbreviated in the text as Tg-*nes*.*cre* or Tg.

## ACKNOWLEDGMENTS

This work was supported by the Isabella and Marcus Paediatric Brainstem Tumour Fund, Victoria, Australia, and the Brainchild Foundation, Queensland, Australia. We thank Jason Cain for providing the Tg-*nes*.*cre* transgenic line.

